# Label-guided seed-chain-extend alignment on annotated De Bruijn graphs

**DOI:** 10.1101/2022.11.04.514718

**Authors:** Harun Mustafa, Mikhail Karasikov, Nika Mansouri Ghiasi, Gunnar Rätsch, André Kahles

## Abstract

Exponential growth in sequencing databases has motivated scalable De Bruijn graph-based (DBG) indexing for searching these data, using annotations to label nodes with sample IDs. Low-depth sequencing samples correspond to fragmented subgraphs, complicating finding the long contiguous walks required for alignment queries. Aligners that target single-labelled subgraphs reduce alignment lengths due to fragmentation, leading to low recall for long reads. While some (e.g., label-free) aligners partially overcome fragmentation by combining information from multiple samples, biologically-irrelevant combinations in such approaches can inflate the search space or reduce accuracy.

We introduce a new scoring model, *<u>m</u>ulti-label <u>a</u>lignment* (MLA), for annotated DBGs. MLA leverages two new operations: To promote biologically-relevant sample combinations, *Label Change* incorporates more informative global sample similarity into local scores. To improve connectivity, *Node Length Change* dynamically adjusts the DBG node length during traversal. Our fast, approximate, yet accurate MLA implementation has two key steps: a single-label seed-<u>c</u>hain-extend <u>a</u>ligner (*SCA*) and a <u>m</u>ulti-label <u>c</u>hainer (*MLC*). *SCA* uses a traditional scoring model adapting recent chaining improvements to assembly graphs and provides a curated pool of alignments. *MLC* extracts seed anchors from *SCA*’s alignments, produces multi-label chains using MLA scoring, then finally forms multi-label alignments. We show via substantial improvements in taxonomic classification accuracy that MLA produces biologically-relevant alignments, decreasing average weighted UniFrac errors by 63.1–66.8% and covering 45.5–47.4% (median) more long-read query characters than state-of-the-art aligners. MLA’s runtimes are competitive with label-combining alignment and substantially faster than single-label alignment.

## 1 Introduction

Sequencing databases are growing exponentially in size [1]. In recent years, *sequence graphs*, in particular *De Bruijn graphs* (DBGs), have become increasingly prominent models for representing and indexing large collections of sequencing data [2], enabling improvements in both the scale and accuracy of many biological analysis tasks, including genotyping [3, 4], variant calling [5, 6], and sequence search [3, 7].

Established search methods are designed for databases of assembled genomes [8]. However, a large fraction of sequencing data deposited in archives like the Sequence Read Archive (SRA) or the European Nucleotide Archive (ENA) have not yet been assembled [9]. This is because assembly requires expensive compute and human labour resources, done by first representing the overlaps between reads as an *assembly graph* (e.g., a DBG), then extensively cleaning the graph (often requiring manual intervention), and finally assembling contiguous sequences (*contigs*) by graph traversal [10]. Since genome assembly strives for long, high-quality contigs [11], the cleaning may discard a significant amount of signal from the sample [12, 13] with no guarantee that there will be no misassemblies among the final contigs [11].

To avoid these signal reduction and misassembly issues, a common way to compress and index a collection of unassembled read sets for search queries is to first construct and only lightly clean an assembly graph for each read set, then merge the graphs into a *joint assembly graph* [7, 14]. *For indexing diverse sequencing data sets, light cleaning is still crucial to reduce the accumulation of noise when indexing diverse collections containing hundreds of thousands of samples [7]*. *DBG-based indexing tools encode metadata as graph annotations*, a key-value store associating each node with one or more metadata tracks, such as sample *labels* [7, 15–23]. *Similar to these previous works, we use accession IDs as labels to associate nodes back to their original database entries*.

*A key search task on these indexes is sequence-to-graph alignment*, a generalisation of pairwise sequence-to-sequence alignment (i.e., computing the maximum similarity *score* between a *query* and a *target* sequence). In this setting, the target sequences are the spellings of contiguous walks (§2.1) on the sequence graph [24–34]. Many of these tools follow a three-step *seed-chain-extend* search paradigm (§2.2). This involves extracting and *anchoring* query *seeds* to the graph, using a *co-linear chaining* algorithm to construct anchor chains, and then *extending* the chains via a search in the graph to form alignments.

### 1.1 Challenges when aligning to annotated De Bruijn graphs

A large proportion of read sets in the SRA are sequenced at low depth^8^, producing heavily fragmented assembly graphs [13]. For metagenomics samples in particular, the constituent organisms are often sequenced at or below 1*×* coverage [35]. Although the light cleaning applied to assembly graphs ensures that exact seed matches can be found [7], the long contiguous walks required for high-scoring (i.e., high-precision) alignments are often not present because of high graph fragmentation. This results in low recall, particularly for long reads, because the short alignments that can be found are not reported to maintain search precision.

Current alignment approaches for sequence graphs have limited support for fragmented graphs. The first approach is *label-free* alignment, which ignores annotations during alignment. When applied to an annotated DBG representing a diverse cohort, these tools [21, 25, 26, 28, 29, 36, 37] can meander search through a large search space inflated by biologically-irrelevant sample combinations [3, 15]. For this reason, these tools primarily target single-species pangenomes that often satisfy the assumption that all walks are biologically relevant. If this assumption holds, then these methods can overcome fragmentation in an individual sample’s assembly graph by combining sequence information from multiple samples since such contiguous walks are present in the joint graph. A second approach aligns queries to walks where all nodes in the walk share one (*single-label*) or more (*label-consistent*) common label(s) [24, 38]. These tools suffer from low recall on fragmented assembly graphs, and so, this property also applies to joint annotated graphs because sample combinations cannot compensate for fragmentation. However, these tools do not suffer from search space inflation because all walks are biologically relevant. This is why these tools are applied to high-quality contiguous graphs indexing reference genomes or high-depth sequencing samples [24, 38]. A third, recently-emerging intermediate approach is *haplotype-aware* alignment, which either aligns in a label-free fashion and scores recombinations afterwards [39] or combines samples to a restricted degree by penalising each combination during alignment [40, 41]. These alignment strategies do not consider similarity (hence, biological relevance) when scoring a sample change and have so far only been applied to single-species pangenomes.

A property shared by all of these approaches is that a discontinuity in an individual assembly graph that does not overlap with another assembly graph will propagate to the joint DBG, limiting the joint graph’s contiguity. This stems from the approaches’ shared definition of alignment: the alignment target must be a contiguous walk.

### 1.2 Contribution: Label-guided sequence-to-DBG alignment

Our goal in this work is to develop an alignment approach that can produce long accurate alignments to collections of low-depth sequencing samples represented by fragmented annotated DBGs. To this end, we propose a new alignment strategy called *<u>m</u>ulti-label <u>a</u>lignment* (MLA) designed for annotated DBGs. The strategy extends alignment scoring models with two key new operations: *(i) Label Change* and *(ii) Node Length Change*. The label change operation penalizes traversals that change from one sample to a dissimilar sample, thus enhancing local single-character similarity scores with more informative global similarity. The node length change operation dynamically adjusts the node length, thus using shorter-length nodes as proxies for missing nodes to locally improve graph connectivity (i.e., reducing fragmentation).

Efficiently implementing these operations must overcome several computational challenges. First, efficient anchor chaining relies on having a small number of anchors per query [42], an assumption that is easily violated in a multi-label setting. Second, sequence-to-graph alignment decision problems (i.e., does there exist an alignment) that allow for DBG edits is shown to be 𝒩𝒫-complete [43], necessitating heuristics to prevent excessive use of node length change operations.

To address these challenges, we implement MLA in a fast, approximate, yet accurate two-step approach: The first step is a new single-label seed-chain-extend aligner for annotated DBGs called *SCA* (Fig 1.1-3), meant to reduce the computational burden of multi-label chaining by first performing single-label chaining with a traditional scoring model and extending the top chains among all labels to provide a preliminary alignment pool. *SCA*, adapts recent improvements in chaining to a DBG setting. The second step is a multi-label chainer called *MLC* (Fig 1.4) that incorporates our novel MLA scoring operations into its chain scoring. It extracts anchors from the alignments provided by *SCA*, resulting in a much smaller curated anchor set on which we apply our more expensive operations. It then performs multi-label chaining on these anchors and stitches the multi-label chains into alignments using fragments from *SCA*’s alignments.

**Fig. 1.**
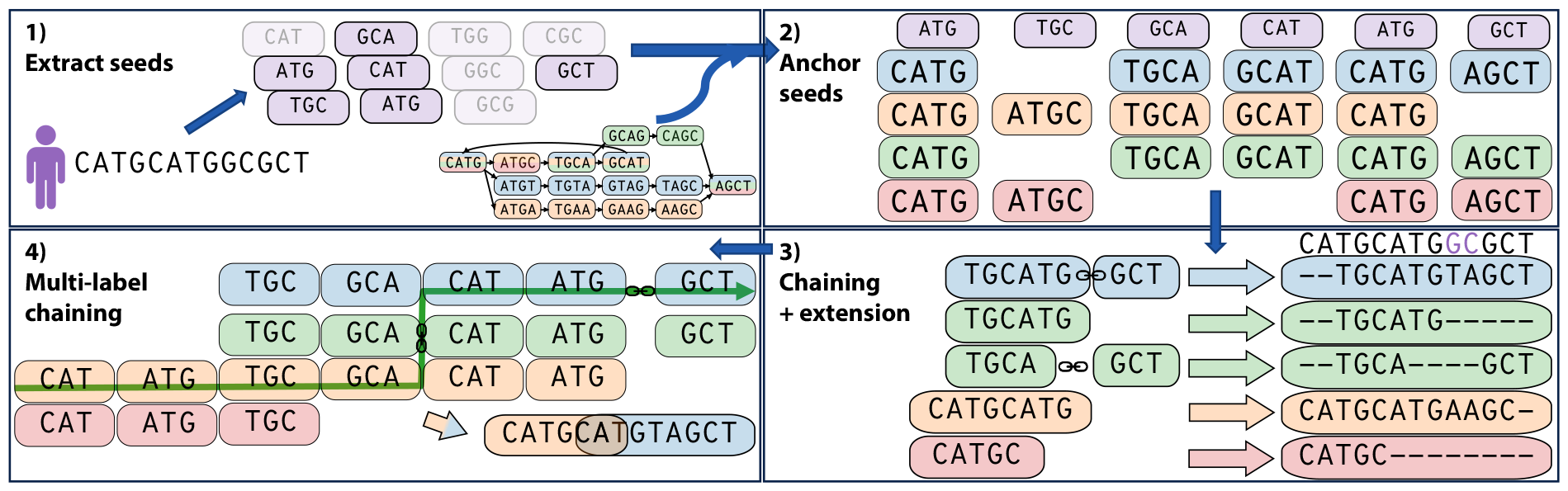
Computing multi-label alignments (MLAs) of a query sequence to an annotated De Bruijn graph. Each colour represents a label in the graph annotation, with some nodes having multiple labels. We first *1)* extract seeds (shown in purple, shaded seeds have no match) of length *l* ≤ *k* from the query sequence (in this example, *k* = 4, *l* = 3, and the query is CATGCATGGCGCT) and *2)* anchor the seeds to the graph, where each anchor matches a seed to a node and a label (each column represents the node-label pairs to which a seed matches). Then, we *3)* construct single-label chains and extend them along single-label walks into alignments using *SCA* (the purple characters in the query indicate mismatched characters). We then *4)* extract anchors from this alignment pool and form multi-label chains using *MLC* (indicated by the green line). We connect anchors using single-label alignment segments to form MLAs.

Overall, we show in this work how our fast approximate implementation of MLA produces substantially longer and more accurate alignments compared to state-of-the-art aligners.

## 2 Preliminaries and Background

### 2.1 Notation and Definitions

A *string* is a finite sequence of characters drawn from an alphabet *Σ. Σ*^*k*^ denotes the set of all strings of length *k* (*k-mers*). For a string *s* = *s*[1]*s*[2] *… s*[*l*] of length |*s*| = *l*, with indices 1 ≤ *i* ≤ *j* ≤ *l*, we denote a substring by *s*[*i* : *j*] := *s*[*i*] *… s*[*j*]. The set of all *k*-mers extracted from a string set *S* is denoted by 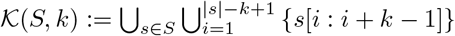.

A *node-centric De Bruijn graph* (DBG) of order *k* representing *S* has the nodes *V* := 𝒦(*S, k*) and implicit edges *E*(*V*) := *{*(*v*_1_, *v*_2_) *∈ V* ^2^ : *v*_1_[2 : *k*] = *v*_2_[1 : *k* − 1]*}* [44]. The *spelling* of a walk *W* = (*v*_1_, …, *v*_*m*_) on a DBG is the string *T* = *v*_1_*v*_2_[*k*] *… v*_*m*_[*k*]. An *annotated DBG* has an auxiliary set of string labels ℒ and an annotation 𝒜 : *V* → 2^ℒ^ associating each node with a label set. *W* is *label consistent* if 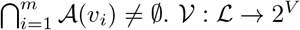 fetches all nodes with a given label.

We denote a query string by *Q ∈ Σ*^|*Q*|^. A *sequence-to-graph alignment* of *Q* to the *target string T* along *W* is a tuple *a* = (*Q*_*a*_, *T*_*a*_, *E*_*a*_, *W*), where *Q*_*a*_ is a substring of *Q, T*_*a*_ is a substring of *T*, and *E*_*a*_ is a sequence of *edit operations* transforming *T*_*a*_ to *Q*_*a*_. These operations are in *{*match, mismatch, insertion open, insertion extension, deletion open, deletion extension*}*. Each operation has a *score*, denoted by *Δ*_=_, Δ≠, *Δ*_*IO*_, *Δ*_*IE*_, *Δ*_*DO*_, and *Δ*_*DE*_, respectively. Only *Δ*_=_ is positive, all other scores are negative. The score of *a*, denoted by *Δ*_*S*_(*a*), is the sum of all edit operation scores, where a higher score indicates greater similarity. *a* is label consistent if *W* is label consistent. Given two alignments *a*_1_, *a*_2_ with respective substrings *Q*[*i*_1_ : *i*_1_ +*l*_1_ −1], *Q*[*i*_2_ : *i*_2_ +*l*_2_ −1] s.t. *i*_1_ *< i*_2_, we define overlap(*a*_1_, *a*_2_) := min*{l*_2_, *i*_1_ + *l*_1_ − *i*_2_*}. a*_1_ and *a*_2_ *overlap* if overlap(*a*_1_, *a*_2_) *>* 0 and are *disjoined* by a *gap* of length −overlap(*a*_1_, *a*_2_) otherwise. For a gap length *l ∈* ℕ^+^, we denote the scoring model’s *gap penalty* by *Δ*_*G*_(*l*). In this work, we assume affine scoring (e.g., *Δ*_*G*_(*l*) := *Δ*_*IO*_ + (*l* − 1)*Δ*_*IE*_ for an insertion).

### 2.2 Sequence-to-graph alignment with seed-extend-chain

Modern approximate aligners predominantly use the *seed-chain-extend* paradigm involving *(i) seed anchoring, (ii) co-linear chaining*, and *(iii) anchor extension* [45]. A *seed* is a query substring while an *anchor* is a tuple of a seed and a graph walk spelling a superstring of the seed. A *chain* is a sequence of anchors s.t. any two consecutive anchors are in order in both the query and target, meaning that there exists a walk connecting their nodes [46]. Some works determine a traversal distance between nodes on-the-fly by traversing local neighbourhoods around nodes [29, 30]. More efficient strategies require a decomposition of the graph, typically into subgraphs [28, 47] or a path/walk cover [31–33, 46]. After chaining, anchor extension searches the graph forwards and backwards from the ends of each anchor to find high-scoring walks.

## 3 Methods

### 3.1 General alignment workflow

Given a query *Q*, we find and anchor seeds, compute label-consistent alignments using *SCA*, then chain these alignments together into MLAs using *MLC*. First, given a user-set seed length *l* ≤ *k*, we extract *l*-mer seeds from *Q* (Fig. 1.1) and anchor them by fetching all graph nodes with matching *l*-length suffixes and their associated labels (Fig. 1.2). An anchor *α*_*ilvℓ*_ is a tuple in *A* := ℕ *×* ℕ *× V × L* anchoring the seed *Q*[*i* : *i* + *l* − 1] to a node *v*, with an associated label *ℓ ∈ 𝒜*(*v*). Afterwards, we find the top-scoring single-label chains (Fig. 1.3, §3.4). We extend each chain into a single-label alignment by using global alignment to connect consecutive anchors and ends-free extension from the first and last anchor in the chain. Given the resulting alignment pool, we extract all *l*-mer anchors and compute multi-label chains (Fig. 1.4, §3.5), incorporating label change (*Δ*_*LC*_) and node length change operations (*Δ*_*L*_). We construct MLAs from the top multi-label chains using segments from the label-consistent alignments.

### 3.2 Deriving a Scoring Model for Novel Alignment Operations

We now detail the scoring model for our new operations for sequence-to-graph alignment: *Δ*_*LC*_ and *Δ*_*L*_. For all constants defined in this section, refer to §4.3 for the values we use in our experiments. We define and extend two probabilistic graphical models for sequence-to-sequence alignment: a *target model* and a *null model*, based on the probabilistic models presented by [48]. Briefly, a model represents all possible mutations of an underlying target sequence, where a walk in a model emits edit operations that generate a query sequence (detailed in Supp. §2.2). The score of any edit operation is the log probability ratio of the operation’s corresponding transition probabilities in the target and the null model. The alignment score is the sum of these log ratios (i.e., the query’s log-likelihood ratio). These sequence-to-sequence alignment models trivially induce analogous models for sequence-to-graph alignment by treating the spelling of each walk in a sequence graph as a separate target sequence and, hence, a pair of target and null graphical models.

#### Deriving and Computing Label-change Scores

For simplicity, suppose that we traverse along an edge (*v*_1_, *v*_2_) s.t. 𝒜(*v*_1_) = *{ℓ*_1_*}* and 𝒜(*v*_2_) = *{ℓ*_2_*}*, where *ℓ*_1_ *≠ ℓ*_2_. We denote this *label change* from *ℓ*_1_ to *ℓ*_2_ as *ℓ*_1_ → *ℓ*_2_ and score this event using the probability that *v*_2_ has label *ℓ*_1_ conditioned on *v*_2_ having the label *ℓ*_2_. Intuitively, this is the probability that the *k*-mer *v*_2_ observed in the sample with label *ℓ*_2_ is also present in the sample with label *ℓ*_1_, but was not observed (Fig. 2). To formulate this precisely, we extend our alignment models so that each graph-traversing operation emits a label change with transition probability Pr(*ℓ*_1_ → *ℓ*_2_). Thus, the *label-change score* is

**Fig. 2.**
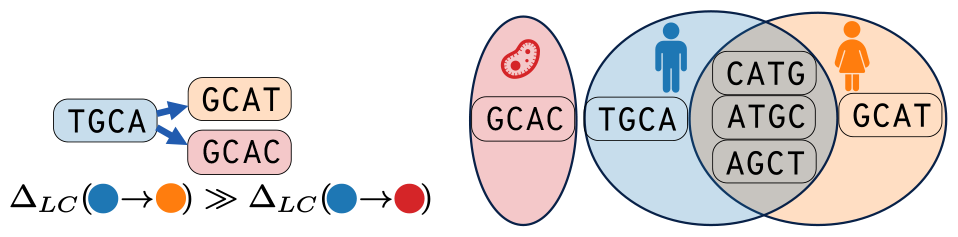
Label change scores measure sample similarity. *Δ*_*LC*_ measures the probability that a node with an orange label is also present in the blue sample but was not observed in that sample. We score a change to the orange label much higher because of the large overlap between the orange and blue *k*-mer sets.

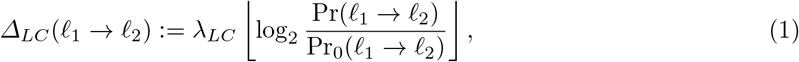

where *λ*_*LC*_ is a user-set scaling constant. For the null model, we assume no relationship between *ℓ*_1_ and *ℓ*_2_, so Pr_0_(*ℓ*_1_ → *ℓ*_2_) := Pr(*ℓ*_2_), where Pr(*ℓ*) is the empirical probability of a label 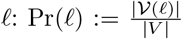. For this new model to be reducible to a label-free setting, we require that *Δ*_*LC*_(*ℓ* → *ℓ*) = 0 (i.e., only score if the label changes) and that *Δ*_*LC*_(*ℓ*_1_ → *ℓ*_2_) ≤ 0 (i.e., no label change increases the score). We satisfy these requirements if Pr(*ℓ*_1_ → *ℓ*_2_) := Pr(𝒱(*ℓ*_1_) ∩ *𝒱*(*ℓ*_2_)), which simplifies Equation 1 to

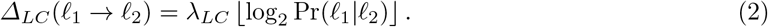

Due to the log in the definition of *Δ*_*LC*_ and the use of integers for alignment scores, we only require order-of-magnitude accuracy when computing Pr(*ℓ*_1_|*ℓ*_2_). So, we approximate Pr(*ℓ*_1_|*ℓ*_2_) using HyperLogLog++ counters [49] of 𝒱(*ℓ*_1_) and 𝒱(*ℓ*_2_), respectively, and the precomputed values |*𝒱*(*ℓ*_1_)| and |*𝒱*(*ℓ*_2_)|.

#### Deriving Penalties for Node Length Changes

Since low sequencing depth can produce disconnected graphs, one way to compensate for this is to allow for node insertions into the graph during the search. In this section, we describe how we use dynamic changes of the underlying DBG’s order *k* during alignment as a more tractable proxy for inserting nodes.

Although *MLC* only utilises node length change operations during chaining, we nonetheless incorporate this operation into our scoring model to derive its scoring function. For this, we switch our graph from a DBG of fixed order *k* to a variable-order DBG of maximum order *k* [50].

Given a node *v* of length *k* with spelling *s* and a suffix length 1 ≤ *l < k*, a node length change is the traversal from *v* to the node with spelling *s*[*k* − *l* + 1 : *k*]. In our model, nodes with length *l* are proxies for missing *k*-mers (Fig. 3). However, we need to define penalties for changing the node length to prevent degenerate cases, such as searches along walks where every node has a short length (e.g., 1 or 2) that exist for every sequence over the alphabet.

**Fig. 3.**
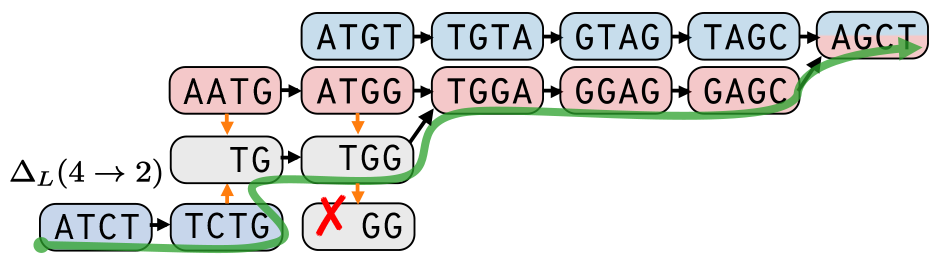
Traversal in a variable-order DBG overcomes graph disconnects to spell the sequence. ATCTGGAGCT. Traversing from `TCTG` to its suffix `TG` allows for traversal to `TGGA`, so the red nodes compensate for the disconnect in the blue subgraph. In MLA, nodes of length *l < k* act as proxies for *k*-mers. In this graph, `TGG` stands in for `CTGG`. In our model, nodes of length *< k* may only traverse to longer nodes (e.g., `TGG` cannot traverse to `GG`). The green line represents the path taken to spell the sequence above and the orange arrows represent node length-changing traversals.

To avoid this case, and to ensure that the model reduces to standard sequence-to-graph alignment on fixed-*k* DBGs, our scoring model does not penalise traversals that increase the node length by 1 or maintain the node length at *k*. Otherwise, we penalise traversal from a node of length *k* to one of length 1 ≤ *l < k* with a score *Δ*_*L*_(*k* → *l*) and disallow all other node length-changing traversals. Consolidating these rules,

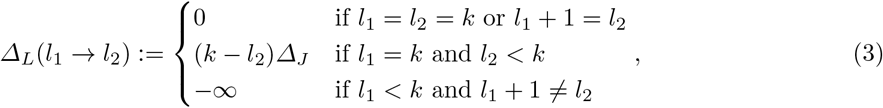

where *Δ*_*J*_ *<* 0 is a user-set constant.

### 3.3 Local co-linear chaining on assembly DBGs

We now describe the modular chaining algorithm used by both *SCA* and *MLC*. Given anchors sorted by increasing end position (i.e., *i* + *l*), we perform local chaining using Algorithm 1, based on the more practical alternative algorithm implementation by [42]^9^. Their algorithm minimises a non-negative chain cost rather than maximising an integer score, so we use the equations by [52] to convert scores into costs when evaluating our termination condition. This *forward pass* computes optimal chaining scores. We reconstruct chains by *backtracking*, ensuring that we incorporate each anchor into, at most, one chain.

The helper function `connect` : *A × A* → ℤ approximates the score of a global alignment connecting anchor *α*_1_ to *α*_2_, with different implementations for *SCA* and *MLC*. Note that this score does not include *α*_1_ since its score is already included in the score vector *Δ*_*S*_ when computing updates.

#### Algorithm 1

Computing local co-linear chaining scores.

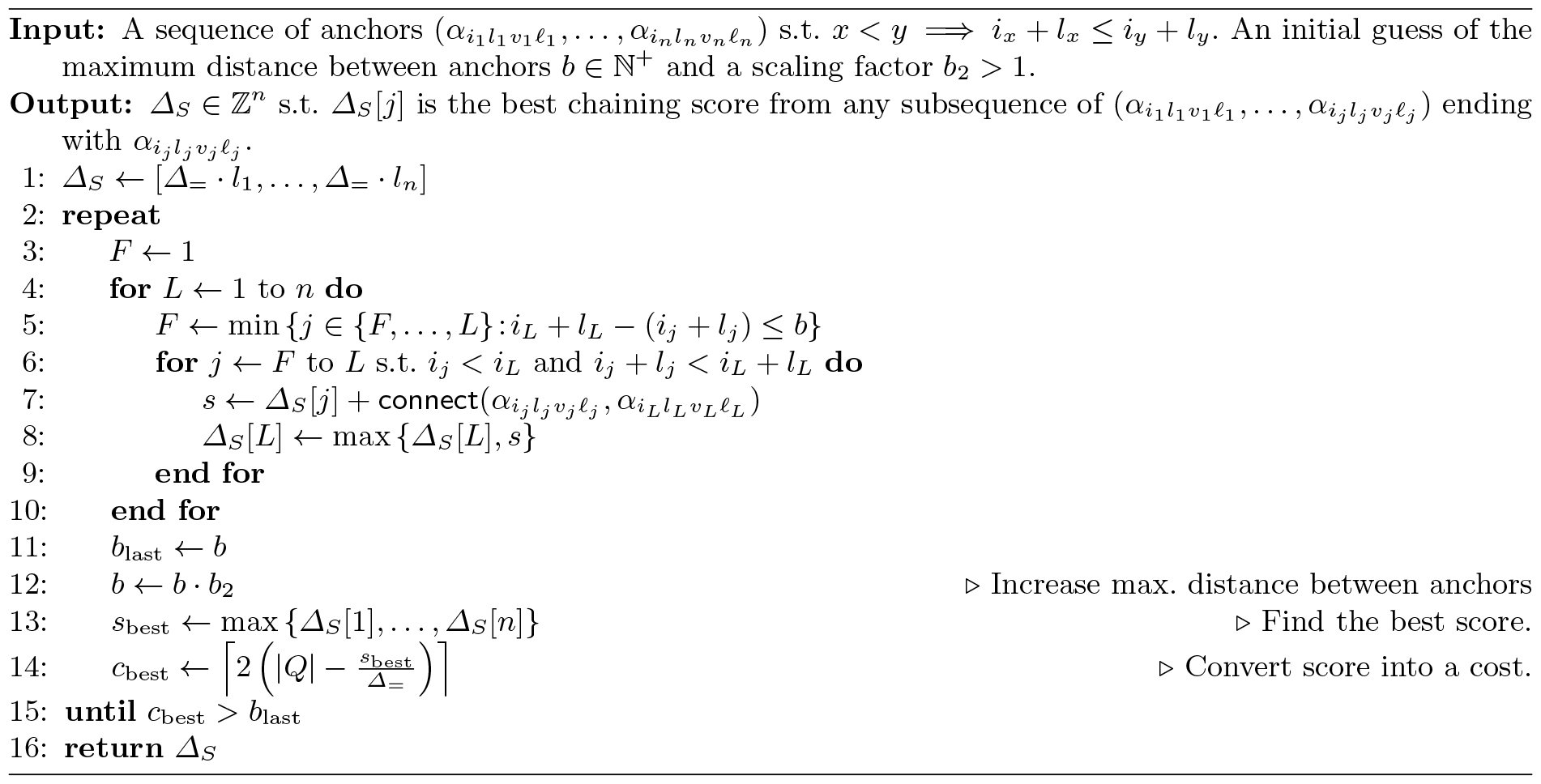

### 3.4 *SCA*: Single-label seed-chain-extend alignment

*SCA* implements co-linear chaining and anchor extension algorithms for label-consistent alignment. After seed anchoring, we merge the anchors into maximal unique matches (MUMs). We then group the anchors by label and perform separate forward passes of the chaining algorithm for each group. Afterwards, we select the top *ρ*|*Q*| chains (with ties) among all labels and backtrack to construct these chains. *ρ* is a user-set chain density (§4.3).

To compute the connection score between two anchors *α*_1_ and *α*_2_, we sum a match score for the additional characters introduced by *α*_2_ to a gap penalty based on the absolute difference between *(i)* the difference of *α*_1_ and *α*_2_’s seed end positions in the query, and *(ii)* the traversal distance between their nodes in the graph (Supp. Algorithm 1). To quickly estimate a traversal distance between two nodes along a label-consistent walk, we use *walk covers* of the subgraphs representing each label. A walk cover is a set of walks s.t. each node is visited at least once. So, a traversal distance is known if there exists a walk from *α*_1_ to *α*_2_ in the cover.

For each chain, we connect consecutive anchors using global alignment, then extend the first anchor backwards and the last anchor forwards using ends-free extension. For global alignment, we use *TCG-Aligner* [15] to ensure that each connecting walk is represented by the cover. For ends-free extension, we modify *TCG-Aligner*’s extender to restrict traversal to label-consistent walks. To further reduce the number of extensions, we discard chains that overlap with already-completed alignments. The result is a pool of label-consistent alignments.

### 3.5 *MLC*: Multi-label co-linear chaining

Given a pool of label-consistent alignments, we extract anchors from these alignments and construct multi-label chains (Fig. 1.4). Unlike the co-linear chaining method in *SCA*, we have access to global alignment scores for connecting any in-order pair of anchors extracted from the same label-consistent alignment. We leverage this to define an anchor connection score that ensures that *MLC*’s chaining scores equal the final MLA scores.

One property of *MLC*’s scoring is that, given a chain of anchors from the same alignment and label, we only need the anchors at the beginning and end of the chain to compute the chain score. So, as a preprocessing step, we discard any anchor 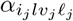 if it is the only anchor at position *i*_*j*_ and if another anchor 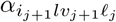 exists from the same alignment.

After the forward pass, we reconstruct the highest-scoring chains that cover each label from the pool of label-consistent alignments. We construct an MLA from each chain by connecting consecutive anchors using alignment segments from the pool.

#### Multi-label anchor connection scoring

We now detail *MLC*’s anchor connection scoring (Supp. Algorithm 2). Suppose we have anchors 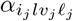 and 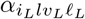 from alignments *a*_*x*_ and *a*_*y*_, respectively. We consider three cases for the update score: *(i)* extending along the same alignment (i.e., *x* = *y*), *(ii)* connecting disjoint alignments (i.e., overlap(*a*_*x*_, *a*_*y*_) ≤ 0), and *(iii)* connecting overlapping alignments (i.e., overlap(*a*_*x*_, *a*_*y*_) *>* 0).

1. If *x* = *y*, then the anchor connection score is the score of *a*_*y*_’s segment up to the longer query substring ending at *i*_*L*_ + *l* (i.e., *Δ*_*S*_(*a*_*y*_[: *i*_*L*_ + *l*]) − *Δ*_*S*_(*a*_*y*_[: *i*_*j*_ + *l*])).
2. If *a*_*x*_ and *a*_*y*_ cover disjoint regions of *Q*, then the connection score includes the sum of *a*_*x*_’s remaining score, the score of *a*_*y*_’s segment up until *i*_*L*_ + *l*, the label change score, and the gap penalty for inserting extra query characters:

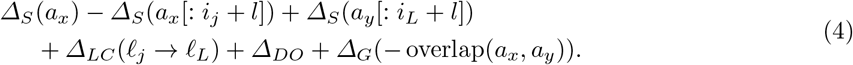

To differentiate this case from *(iii)*, we insert a sentinel $ character into the target spelling with an incurred *Δ*_*DO*_ score.
3. If *a*_*x*_ and *a*_*y*_ cover overlapping regions of *Q*, assume that the current alignment segment of *a*_*x*_ up until *i*_*j*_ + *l* is of length *≥ k* and that |*a*_*y*_[*i*_*L*_ :]| *≥ k* (meant to avoid ambiguity when spelling a walk). We also only consider overlaps of length *< k* − 1 (i.e., one cannot traverse from *a*_*x*_ to *a*_*y*_ with a single step without modifying the graph) since we assume that longer overlaps would have been discovered otherwise via normal graph traversal during anchor extension. We use a two-step procedure to try to connect the two anchors. First, we try to find an intermediate anchor 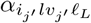 from *a*_*y*_ s.t. *i*_*j*_ = *i*_*j*_*′* and hop to that node. Then we continue the traversal along *a*_*y*_ towards *v*_*j*_*′* to complete the connection. If such a node exists, the connection score is

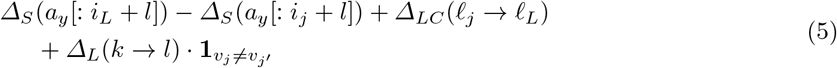

Using Fig. 3 as an example, one such intermediate anchor is the node `AATG` sharing a 2-mer suffix with the node `TCTG`.

Note that this score does not depend on the length of *v*_*j*_ and *v*_*j*_*′* ‘s longest common suffix if *v*_*j*_*≠ v*_*j*_*′*. We motivate this design choice with the following observation: If there is a single character mismatch between position *k* − *l*^*′*^ + 1 (where *l*^*′*^ need not equal the seed length *l*) in a node *v*’s spelling and position *k* − *l*^*′*^ in another node *v*^*′*^’s spelling that prevents the nodes from being adjacent (e.g., *v* spells `GATGC` and *v*^*′*^ spells `ACGCT` for *k* = 5 and *l*^*′*^ = 3), then we require *l*^*′*^ node insertions to create a path to *v*^*′*^ from a predecessor of *v*, despite the ground truth that it originated from a single substitution (for our example, given appropriate characters *c, d ∈ Σ*, these new nodes would spell *cd*`GAC`, *d*`GACG`, and `GACGC`). Thus, defining the connection score as a function of *k* − *l*^*′*^ instead of *k* − *l* would induce unequal scores for each value of *l*^*′*^, even though all of these edit events are equally likely.

### 3.6 Time and Space Complexity

Suppose we have *n* sorted anchors (incurring a worst-case complexity of 𝒪(*n* log *n*) if the seeder does not produce sorted anchors). Let *C* denote the time to execute a `connect` function. If *n* ≤ |*Q*|, then Algorithm 1 has an average-case time complexity in 𝒪(*c*^*\**^|*Q*|*C*), where *c*^*\**^ is the minimum chain cost [42]. If *n >* |*Q*|, then we fall back to the chaining algorithm from minimap2 [51] with a worst-case time complexity in 𝒪(*bnC*) for the forward pass, where *b* is a user-set bandwidth parameter. Although we perform *ρ*|*Q*| backtracking procedures, since each anchor is incorporated into at most one chain, we terminate a procedure as soon as we reach an already-chained anchor. So, backtracking takes worst-case 𝒪(*n*) time. Alongside the *n* anchors, we store *Θ*(1) information per anchor for the chaining scores and backtracking information, so the total space complexity is in *Θ*(*n*).

Consider *SCA*’s `connect`. Suppose that *Q* maps to *L* labels, with corresponding anchor counts *n*_1_, …, *n*_*L*_, walk cover sizes *W*_1_, …, *W*_*L*_, and optimal chaining costs 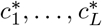. Since MUMs generally satisfy *n*_*i*_ ≤ |*Q*| [42], the average-case time complexity of *SCA*’s forward passes is 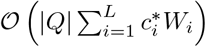 After extension, we extract *m* anchors from the resulting alignments.

Considering *MLC*, it is clear that *C*_MLA_ *∈ 𝒪*(1). Since we chain anchors from multiple labels in a single forward pass, we can no longer assume that *m* ≤ |*Q*| will generally hold. Thus, MLA chaining has an average-case time complexity in 𝒪(*c*^*\**^|*Q*|) when *m* ≤ |*Q*| and worst-case time complexity in 𝒪(*bm*) otherwise. Splicing alignment segments to convert a multi-label chain into an MLA spelling *T* runs in worst-case 𝒪(|*T* |) time.

## 4 Evaluation Methodology

We compare our methods to *GraphAligner* [28], a state-of-the-art tool for traditional sequence-to-graph alignment to DBGs, and *PLAST* [24], a BLAST-like tool for label-consistent alignment to annotated DBGs. We implement two additional baselines: *SCA+FixedMLC100* implements multi-label chaining with *Δ*_*LC*_ = −100, while *SCA+MLC(no NLC)* implements our *Δ*_*LC*_ but disallows node length changes. We evaluate all tools on a simulated joint assembly graph indexing 3042 fungi genomes from GenBank [53].

### 4.1 Simulating assembly graphs and query sequences

From each genome, we simulate a 10*×*-coverage Illumina HiSeq-type read set using *ART* v2.5.8 [54] and construct a cleaned assembly DBG of order *k* = 31 using the procedure in *MetaGraph* [7]. We then merge these graphs into an annotated DBG with nodes labelled by their source accession IDs. To merge the graphs, we compute walk covers for each graph and construct the joint graph from these sequences. We pre-process the graph and generate walks using the procedure described by [33]. To reduce the final representation size, we refrain from maintaining a matrix of traversal distances from each node to each stored walk. Instead, we augment each cover with additional walks spanning the edges discarded during pre-processing. We losslessly encode the walks as coordinate graph annotations [15]. We use the GFA representation of the joint graph to interface with *GraphAligner* and the walk covers to construct the *PLAST* index.

We simulate query reads from the same genomes with a different random seed, using *ART* for Illumina HiSeq-type reads and *pbsim3* [55] for PacBio Sequel CLR-type, PacBio Sequel HiFi-type, and ONT-type reads. We generate HiFi reads from simulated 10-pass Sequel subreads using the PacBio `ccs` tool. We form a query set for each read type by drawing 100 random reads from the pool of reads aggregated from all genomes.

### 4.2 Recall-, coverage- and taxonomy-based measures

Alongside read mapping quality, we measure the accuracy of the retrieved labels w.r.t. the ground-truth genomes using *taxonomic profiles* since our query reads are simulated from known genomes.

Taxonomic classification accuracy is highly dependent on which alignments are reported in a read mapping. So, we first find an appropriate score cut-off to determine which alignments to report for each read. Since our test reads are of different lengths, we divide each score by the length of its corresponding query to get a *relative score*. Given the alignments of a read sorted by decreasing relative score and a cut-off, we select alignments by greedily picking a disjoint subset whose relative scores are above the cut-off. We vary the cut-off from 0.0 to 1.0 in steps of 0.02. For each cut-off, we evaluate the *(i)* recall and the *(ii)* mean taxonomic profile error. We measure taxonomic profile error using the WGSUniFrac error of the profile relative to the ground-truth profile [56], a measure of the fraction of the taxonomic tree traversal distance that differs between the two profiles. For easier interpretation, we define the *UniFrac accuracy* as 1.0 − WGSUniFrac error. Since taxonomic IDs are not available for all strains, we generate a custom taxonomic tree by augmenting the NCBI Taxonomy with a new leaf node for each GenBank accession ID. The ground-truth profile of a read states that its ground-truth accession is 100% abundant, whereas the profiles computed from alignments weight each accession by the fraction of query characters covered by alignments to that accession.

### 4.3 Experimental setup and code availability

We performed all experiments on an AMD EPYC-Rome processor from ETH Zurich’s high-performance compute systems using a single thread and a seed size of *l* = 19, with default parameters for all tools except for the following: For scoring, we set *Δ*_=_ = 1 and *Δ*_*≠*_ = *Δ*_*IO*_ = *Δ*_*IE*_ = *Δ*_*DO*_ = *Δ*_*DE*_ = −1. For *SCA* and *MLC*, we use a chain density of *ρ* = 0.01 and chaining parameters *b* = 400 and *b*_2_ = 4. In *MLC*, we set the label-change score scaling factor to *λ*_*LC*_ = *Δ*_=_ and the node length change scaling factor to *Δ*_*J*_ = *Δ*_*DE*_. We implemented our methods within the GPLv3-licensed *MetaGraph* framework [7] hosted at https://github.com/ratschlab/metagraph. The data and scripts for reproducing our results are available from the `biorxiv` branch at https://github.com/ratschlab/mla/tree/biorxiv.

## 5 Results and Discussion

### 5.1 Assembly graphs are much wider than pangenomes

Our simulated assembly graphs have a median (mean) graph size of 48,660 (122,826) *k*-mers (Supp. Fig. 1). After merging these assembly graphs, the annotated DBG contains 187,662,586 *k*-mers. We observe that unlike the pangenome graphs explored in previous chaining works with widths ranging from 1 to 60 [31– 33], our assembly graphs are much wider (i.e., their minimal walk covers are large), with a median (mean) width of 221 (542.8). These observations corroborate our choice to refrain from encoding a full node-to-walk distance matrix for *SCA*.

### 5.2 Multi-label alignments are substantially longer, computed at competitive execution times

For all read types, we observe that full MLA (*SCA*+*MLC*) produces the longest alignments (Fig. 4), with median read coverage increases of 47.4% for HiFi reads, 46.7% for CLR reads, and 45.5% for ONT reads, respectively, relative to the next-best state-of-the-art aligner. All methods have median coverages of 100% for Illumina reads, with the smallest inter-quartile range observed for MLA. Our baselines *SCA+FixedMLC100* and *SCA+MLC(no NLC)* achieve similar or slightly improved performance relative to *SCA*, but far below MLA. *PLAST* has the highest third quartile coverage for PacBio reads among all tools and higher median coverage on HiFi reads than the most comparable tool *SCA*. However, its execution time is ∼280–1200*×* slower than *SCA* (Fig. 7).

**Fig. 4.**
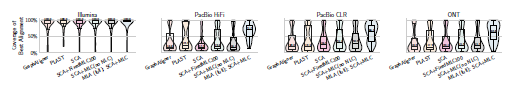
Coverage of each read’s best alignment. Coverage is the percentage of query characters covered by an alignment. *SCA+FixedMLC100* sets *Δ*_*LC*_ = −100 for label changes. *SCA+MLC(no NLC)* uses our *Δ*_*LC*_, but not does allow node length changes.

### 5.3 MLA maintains the best classification accuracy at most read mapping recall levels

Sweeping through different relative score cut-offs, we observe that full MLA has the greatest recall on error-prone reads for all cut-offs and the greatest recall for HiFi reads at cut-offs below 0.5 (Fig. 5). All tools perform similarly on Illumina reads. When comparing taxonomic profiles, MLA has the greatest UniFrac accuracy for long reads at recall values above 20% for HiFi and ONT reads and above 50% for CLR reads (Fig. 6). All tools perform similarly on Illumina reads. For a score cut-off of 0.0, all methods have notable decreases in UniFrac accuracy. The areas under the mean UniFrac Accuracy-Recall curves (AUARCs) for Illumina reads are 0.90 for all methods. For all long-read types, MLA has the highest AUARCs, ranging from 0.85–0.88, compared to *SCA* (0.80–0.83), *SCA+FixedMLC100* (0.82– 0.84), *SCA+MLC(no NLC)* (0.82–0.85), and *PLAST* (0.55–0.68). We infer from these results that MLA improves accuracy by chaining together alignments to related samples.

**Fig. 5.**
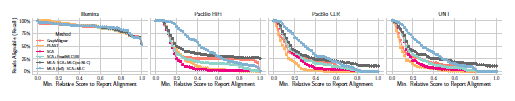
Read mapping recall for different relative score cut-offs. For each relative score (i.e., score scaled by query length) cut-off, the recall is the fraction of reads with an alignment scoring at least at that cut-off.

**Fig. 6.**
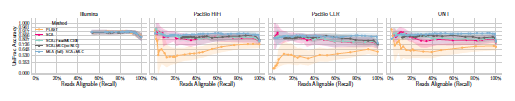
Mean UniFrac accuracy of all mapped reads at different read mapping recall levels. UniFrac accuracy measures the similarity between a read’s taxonomic profile (induced from the labels of all reported alignments) and its ground-truth profile, similar to precision. We estimate means from 1000 bootstrap samples, with shading representing the 95% CI of the mean. Based on our interpretation of WGSUniFrac error values (detailed in Supp. §2.4), each y-axis grid line corresponds to the midpoint value of a taxonomic rank (accession, strain, species, genus, etc.). The y-axis is scaled to better emphasise the upper range of UniFrac accuracy values, corresponding to accuracy at lower taxonomic ranks. We exclude *GraphAligner* since it does not consider labels.

### 5.4 Overall performance

Overall, MLAs are substantially longer than alignments produced by other tools (Fig. 7). We achieve these results while maintaining competitive execution times and the lowest RAM usage. For Illumina reads, *SCA* and *MLC* are substantially faster than all other tools. *GraphAligner* is the fastest tool for long reads, with *SCA* and MLA having comparable execution times.

**Fig. 7.**
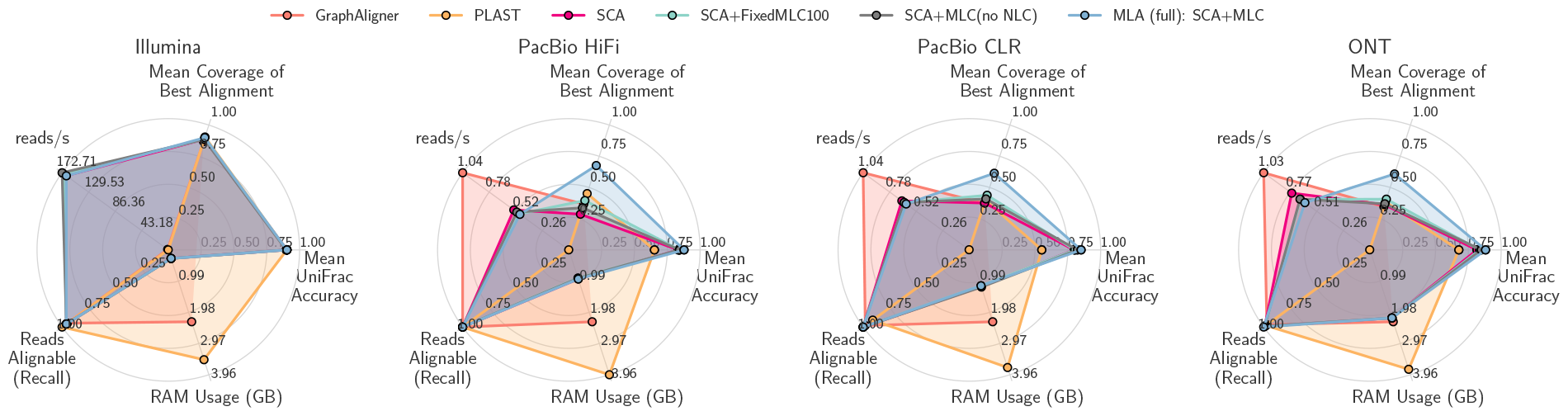
Comparison of performance measures for each alignment tool. See Supp. Tab. 2 for the numbers plotted here.

## 6 Conclusions

In this work, we presented the multi-label alignment (MLA) strategy, a novel alignment scoring model that compensates for disconnects in low-depth assembly graphs by combining sequence information from multiple related samples and improving graph connectivity during alignment. We implement MLA on annotated De Bruijn graphs (DBGs) with a two-step process: *SCA* computes label-consistent alignments and *MLC* computes multi-label alignments from alignments produced by *SCA*.

Since *SCA* does not encode a traversal distance matrix from each node to each walk encoded by the graph annotation, providing such a matrix can potentially increase label-consistent alignment lengths and consequently provide a stronger basis for *MLC*. Despite the large widths of assembly graphs, we expect this matrix to be sparse and, hence, easily compressible.

Another possible extension is merging *SCA* and *MLC* into a single holistic seed-chain-extend procedure. We maintained these two steps in this work to provide a small number of anchors to each chaining run. One can explore how well our current approach approximates this unified approach and how to approximate an ends-free extension incorporating label and node length changes. In this context, there are a few possibilities for implementing the node length change operation into chain scoring. These include using the current procedure of finding intermediate suffix-sharing nodes and a more sensitive, but daunting approach of representing walk covers for all desirable node length values *l < k*.

Although our algorithms are implemented within the *MetaGraph* framework, the concepts from our methods can readily be applied in other pangenome graph frameworks and potentially see more widespread use. These methods make unassembled read sets a more powerful resource for bioinformatics research.

## Acknowledgements

We thank Ragnar Groot Koerkamp, Maximilian Mordig, Mohammed Alser, Mario Stanke, Inanc Birol, and anonymous reviewers for their helpful discussions and feedback. H.M. and M.K. were partially funded as part of Swiss National Research Programme (NRP) 75 “Big Data” by the SNSF grant #407540 167331.

A.K. is partially funded by The LOOP Zurich and the Monique Dornonville de la Cour Foundation to G.R. H.M., M.K., and A.K. were also partially funded by ETH core funding (to G.R.). H.M. is also partially funded by the Personalized Health and Related Technologies (PHRT) Transition Postdoc Fellowship Project #2021/453. We declare no conflicts of interest.

## Footnotes

8 The fungi SRA samples indexed by [7] have a median (mean) *k*-mer multiplicity of 9 (10.5) (Supp. Fig. 2).

9 We fall back to the chaining algorithm from minimap2 [51] with affine gap scoring if there are more than |Q| anchors (§3.6)

