## Supplement for "Label-guided seed-chain-extend alignment on annotated De Bruijn graphs"

### 1 Supplementary Algorithms

**Algorithm 1** connect$(\cdot, \cdot)$ function used by *SCA* to calculate the score for connecting two anchors with the label $\ell$.

**Input:** Anchors $\alpha_{i_j l_j v_j}$ and $\alpha_{i_L l_L v_L}$ with label $\ell$ such that $i_j < i_L$ and $i_j + l_j < i_L + l_L$. A walk cover $\mathcal{W} = \{(v_{11}, \ldots, v_{1n_1}), \ldots, (v_{w1}, \ldots, v_{wn_w})\}$ of the subgraph of all nodes with label $\ell$. Match open score $\Delta_{=} > 0$. Gap penalty function $\Delta_G$.
**Output:** A score $s \in \mathbb{Z}$.

$d_Q \leftarrow i_L + l_L - (i_j + l_j)$
$g_{\text{best}} \leftarrow \infty$
**for all** $(v_1, \ldots, v_n) \in \mathcal{W}$ **do**
    **if** $(v_j, v_L) = (v_x, v_y)$ and $x < y$ **then**
        $g_{\text{best}} \leftarrow \min\{g_{\text{best}}, |y - x - d_Q|\}$
    **end if**
**end for**
**return** $\Delta_{=} \cdot \min\{d_Q, l_L\} + \Delta_G(g_{\text{best}})$

**Algorithm 2** connect$(\cdot, \cdot)$ function used by *MLC* to calculate the score for connecting two anchors of length $l$.

**Input:** Anchors $\alpha_{i_j l v_j \ell_j}$ and $\alpha_{i_L l v_L \ell_L}$ that originate from alignment $a_x$ to label $\ell_j$ and $a_y$ to label $\ell_L$, respectively, such that $i_j < i_L$. A deletion open score $\Delta_{DO} < 0$. Gap penalty function $\Delta_G$.
**Output:** A score $s \in \mathbb{Z}$.

**if** $x = y$ **then**
    **if** $\ell_j = \ell_L$ **then return** $\Delta_S(a_y[: i_L + l]) - \Delta_S(a_y[: i_j + l])$
    **else return** $-\infty$
    **end if**
**end if**
**if** current segment of $a_x$ has length $< k$ or $|a_y[i_L :]| < k$ **then**
    **return** $-\infty$
**end if**
$s \leftarrow \Delta_{LC}(\ell_j \rightarrow \ell_L)$
$o \leftarrow \text{overlap}(a_x, a_y)$
**if** $o \leq 0$ **then**
    **return** $s + \Delta_{DO} + \Delta_G(-o) + \Delta_S(a_x) - \Delta_S(a_x[: i_j + l]) + \Delta_S(a_y[: i_L + l])$
**end if**
**if** $\alpha_{i_{j'} l v_{j'} \ell_L}$ from $a_y$ exists s.t. $i_{j'} = i_j$ **then**
    **if** $v_j \neq v_{j'}$ **then**
        $s \leftarrow s + \Delta_L(k \rightarrow l)$
    **end if**
    **return** $s + \Delta_S(a_y[: i_L + l]) - \Delta_S(a_y[: i_j + l])$
**end if**
**return** $-\infty$

### 2 Supplementary Sections

#### 2.1 Merging single-label anchors into MUMs

We define a *unipath* as a maximal non-branching path in a DBG whose spelling is called a *unitig*. In the initial anchor set, all anchors are of the same length $l$. Then, given two consecutive anchors $\alpha_{i_1 l v_1}$ and $\alpha_{i_2 l_2 v_2}$ (since we may be merging into a previously merged anchor), if $i_1 + l + 1 = i_2 + l_2$, if these are the only anchors with end positions $i_1 + l$ and $i_2 + l_2$, if $(v_1, v_2) \in E$, and if $v_1$ and $v_2$ are in the same unipath, we merge them into a single anchor of length $l_2 + 1$.

#### 2.2 Probabilistic graphical alignment models

Given a starting node in the graph, the first generative model (called the *target model*) generates query sequences with probabilities proportional to their similarity to the spellings of walks that originate at the starting node, assuming that the query sequence is generated by a series of edits to a walk spelling. The second model (called the *null model*) generates both query sequences and graph walks with probabilities only proportional to the sequences' respective lengths, assuming that the two tasks occur independently. Each alignment operation $E$ (defined in Main Section 2.1) corresponds to traversing an edge in the graphical model (called a *transition*), which occurs with a transition probability denoted by $\Pr(E)$ for the target model and $\Pr_0(E)$ for the null model. The graphical model *emits* a query character if $E \in \{=, \neq, IO, IE\}$ and it traverses a graph edge to emit a target character if $E \in \{=, \neq, DO, DE\}$. We define the score of $E$ as

$$\Delta_E := \lambda_E \left\lfloor \log_2 \frac{\Pr(E)}{\Pr_0(E)} \right\rfloor , \qquad (1)$$

where $\lambda_E$ is a user-set scaling constant. Thus, the sum of all operation scores (i.e., the alignment score $\Delta_S$) corresponds to the log-likelihood ratio of the corresponding walk probabilities in the target and null models.

#### 2.3 Assembly graphs have large widths

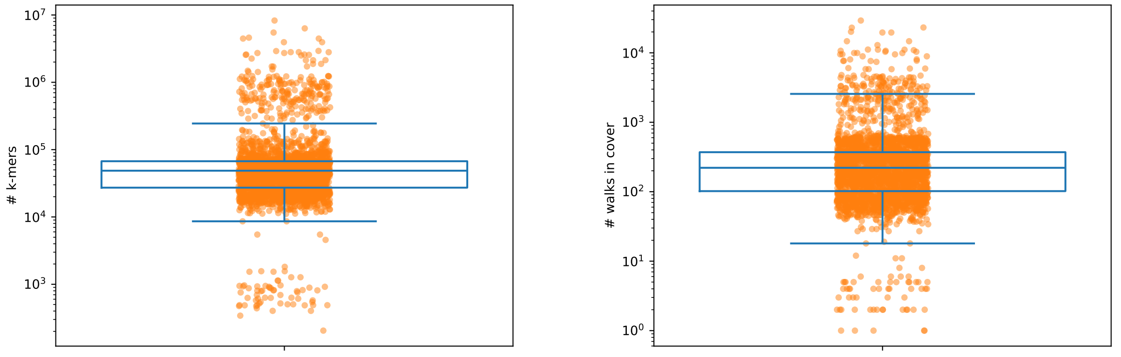


**Fig. 1. Number of $k$-mers (left) and walk cover sizes (right) in the simulated assembly graphs.**.

#### 2.4 Inferring taxonomic ranks from WGSUniFrac values

When computing WGSUniFrac errors, the algorithm assigns a branch length of $x^{-1}$, where $x$ is the distance from the branch to the tree root. We consider nine different ranks: superkingdom, phylum, class, order, family, genus, species, strain, accession, with corresponding branch lengths $\left(\frac{1}{2}, \frac{1}{3}, \frac{1}{4}, \frac{1}{5}, \frac{1}{6}, \frac{1}{7}, \frac{1}{8}, \frac{1}{9}\right)$. If we assume that the taxonomic profile of a read contains a single accession (and by definition, the ground-truth profile has a single accession), then the unweighted WGSUniFrac error for accuracy at level $x$ (meaning a mismatch at rank $x+1$) is $2 \cdot \sum_{i=x+1}^{9} \frac{1}{i}$ since there is an incorrect branch in both profiles. Since the maximum WGSUniFrac error value is ∼5.66, each unweighted error is divided by this constant. Based on this interpretation, we calculate midpoints for WGSUniFrac error and accuracy value ranges corresponding to classification accuracy at least taxonomic rank in Table 1.

**Table 1. Midpoint WGSUniFrac error and accuracy values for matching accuracy at each taxonomic rank**. UniFrac accuracy is $1-$ WGSUniFrac error.

| **Taxonomic Rank** | **WGSUniFrac Error Midpoint** | **UniFrac Accuracy Midpoint** |
|---|---:|---:|
| Accession | 0.0 | 1.0 |
| Strain | 0.039 | 0.961 |
| Species | 0.083 | 0.917 |
| Genus | 0.134 | 0.866 |
| Family | 0.193 | 0.807 |
| Order | 0.264 | 0.736 |
| Class | 0.352 | 0.648 |
| Phylum | 0.470 | 0.530 |
| Superkingdom | 0.647 | 0.353 |
| Root | 1.0 | 0.0 |

#### 3 Supplementary Figures

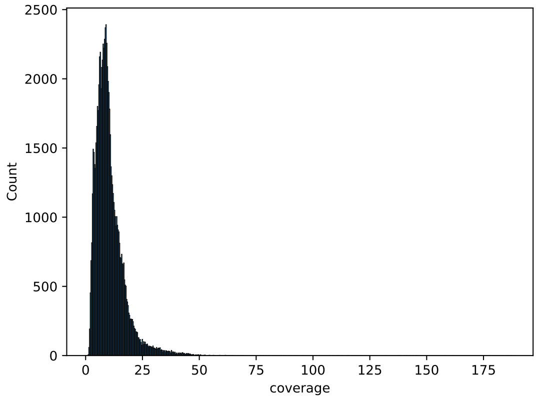


**Fig. 2. $k$-mer count distribution of SRA Fungi samples.**

#### 4 Supplementary Tables

**Table 2. Alignment statistics**

| Platform | Method | Cov. of Best Alignment mean ± std | Recall max | WGSUniFrac Error mean ± std | Time (s) | RAM Usage (GB) |
|---|---|---|---|---|---|---|
| Illumina | GraphAligner | 0.868 ± 0.274 | 0.950 | NaN | 91.740 | 2.281 |
| | PLAST | 0.904 ± 0.200 | 1.000 | 0.147 ± 0.198 | 739.580 | 3.486 |
| | SCA | 0.886 ± 0.247 | 0.960 | 0.119 ± 0.160 | 0.602 | 0.270 |
| | SCA+FixedMLC100 | 0.886 ± 0.247 | 0.960 | 0.119 ± 0.160 | 0.587 | 0.274 |
| | SCA+MLC(no NLC) | 0.886 ± 0.247 | 0.960 | 0.119 ± 0.160 | 0.579 | 0.272 |
| | MLA (full): SCA+MLC | 0.899 ± 0.243 | 0.960 | 0.118 ± 0.161 | 0.603 | 0.270 |
| ONT | GraphAligner | 0.346 ± 0.346 | 0.970 | NaN | 97.300 | 2.281 |
| | PLAST | 0.384 ± 0.362 | 0.980 | 0.289 ± 0.351 | 58362.000 | 3.790 |
| | SCA | 0.362 ± 0.334 | 0.990 | 0.187 ± 0.242 | 132.187 | 2.161 |
| | SCA+FixedMLC100 | 0.406 ± 0.345 | 0.990 | 0.178 ± 0.237 | 148.838 | 2.156 |
| | SCA+MLC(no NLC) | 0.369 ± 0.324 | 0.990 | 0.167 ± 0.211 | 147.848 | 2.160 |
| | MLA (full): SCA+MLC | 0.608 ± 0.283 | 1.000 | 0.139 ± 0.158 | 159.027 | 2.160 |
| PacBio CLR | GraphAligner | 0.373 ± 0.355 | 0.980 | NaN | 96.260 | 2.281 |
| | PLAST | 0.398 ± 0.380 | 0.910 | 0.346 ± 0.400 | 42152.000 | 3.733 |
| | SCA | 0.375 ± 0.309 | 1.000 | 0.231 ± 0.293 | 151.475 | 1.161 |
| | SCA+FixedMLC100 | 0.438 ± 0.340 | 1.000 | 0.227 ± 0.293 | 157.384 | 1.141 |
| | SCA+MLC(no NLC) | 0.408 ± 0.325 | 1.000 | 0.214 ± 0.263 | 156.706 | 1.179 |
| | MLA (full): SCA+MLC | 0.614 ± 0.281 | 1.000 | 0.173 ± 0.195 | 161.772 | 1.138 |
| PacBio HiFi | GraphAligner | 0.368 ± 0.369 | 1.000 | NaN | 96.340 | 2.281 |
| | PLAST | 0.451 ± 0.379 | 1.000 | 0.246 ± 0.308 | 95124.000 | 3.963 |
| | SCA | 0.286 ± 0.263 | 1.000 | 0.185 ± 0.235 | 185.907 | 0.921 |
| | SCA+FixedMLC100 | 0.395 ± 0.342 | 1.000 | 0.182 ± 0.236 | 206.080 | 0.894 |
| | SCA+MLC(no NLC) | 0.337 ± 0.327 | 1.000 | 0.160 ± 0.189 | 196.678 | 0.901 |
| | MLA (full): SCA+MLC | 0.676 ± 0.226 | 1.000 | 0.149 ± 0.172 | 209.100 | 0.924 |
